# Implementation and testing of a biohybrid transition microelectrode array for neural recording and modulation

**DOI:** 10.1101/2023.08.08.552362

**Authors:** Benozir Ahmed, Simon Binder, Saeed Boroomand, Hunter J. Strathman, Michael Hantak, Tate Shepherd, Adrian Ehrenhofer, Huannan Zhang, Jason Shepherd, Florian Solzbacher, Patrick A. Tresco, Steve Blair, Christopher F. Reiche, Yantao Fan

## Abstract

Recent advances in microtissue engineered neural networks have opened avenues for the fabrication of biohybrid intracortical brain implants. To date, however, the experimental validation of these biohybrid implants has been restricted to singularly positioned electrodes, with the functionality confined to optical neuromodulation and neural recording. We present for the first time a biohybrid transition microelectrode array (TMEA) comprising a 4×4 matrix of biohybrid electrodes. These electrodes are expected to project axons into the brain, thereby establishing synaptic connections for electrical readout and excitation. Our device thus offers a promising pathway to leverage organic, endogenous materials in intracortical implants to enhance their biocompatibility as well as their functionality. In particular with respect to functionality, we assume that the spatiotemporal resolution of recorded neural signals may be significantly increased using this technology owing to the size ratios of these biohybrid microelectrodes compared to abiotic ones. Future investigations will therefore focus on the exploitation of the electrode density to further advance brain-computer interface technology.

---

Integral to the restoration of sensory and motor functions as well as the identification of neurological disorders, implanted microelectrode arrays continue to be a key focal point in neuroscientific studies [1–3]. Traditional designs largely utilize rigid silicon microelectrode shafts for both neural stimulation and activity recording [4]. However, the discrepancy between these stiff microelectrodes and the malleable nature of brain tissue can trigger foreign body reactions such as chronic inflammation, bioencapsulation, electrode corrosion, and eventually culminate in signal loss and device failure [5–8]. Methods like parylene C encapsulation, while commonly used, extend the electrode’s mean-time-to-failure without significantly decreasing device stiffness or forestalling eventual bioencapsulation[9–12].

The recent advent of flexible probes has shown promise in addressing these challenges [13]. Our work advances by introducing a novel neural implant that improves on the material incongruity issue by substituting exogenous electrode materials with endogenous ones. At the heart of this implant are microtissue engineered neural networks, a close mimic of brain tissue [14]. With the inclusion of such neural networks into an electrode array, we term this innovative design as a “biohybrid transition microelectrode array (TMEA)”.

Fig. 1 a to 1 c illustrate the functional principle as well as the design of our novel TMEA intracortical neural implant. The polymer microneedles have channels containing a hydrogel material to facilitate neural growth. This microneedle array is attached to a silicon-based chip which carries living neurons in gold-coated microwells (µWells) on the surface. The immobilized neurons are intended to project their neurites through the hydrogel-filled shaft channel and, after implantation, form synaptic connections with the brain’s local neurons. They thus function as a living interlink between the base chip’s electrical recording/stimulating site and the region of interest in the human brain’s cortex. As the cultured neural cells in the µWells are in direct contact with the base chip’s recording sites and at the same time directly connected to the brain’s neural networks, we hypothesize that information will be transported from the deeper brain layers to the surface, where it eventually can be recorded with conventional neuroscience instrumentation equipment. A biohybrid device like this is expected to have a significantly lower mechanical mismatch compared to silicon microneedles of existing designs and, due to its tissue-like nature, show improved shortterm and long-term implant integration. At the same time, it is expected to enhance the potential for large-scale neural recording with high spatiotemporal resolution due to the small size of the neurite-based interlinks into the brain.

**Fig. 1.**
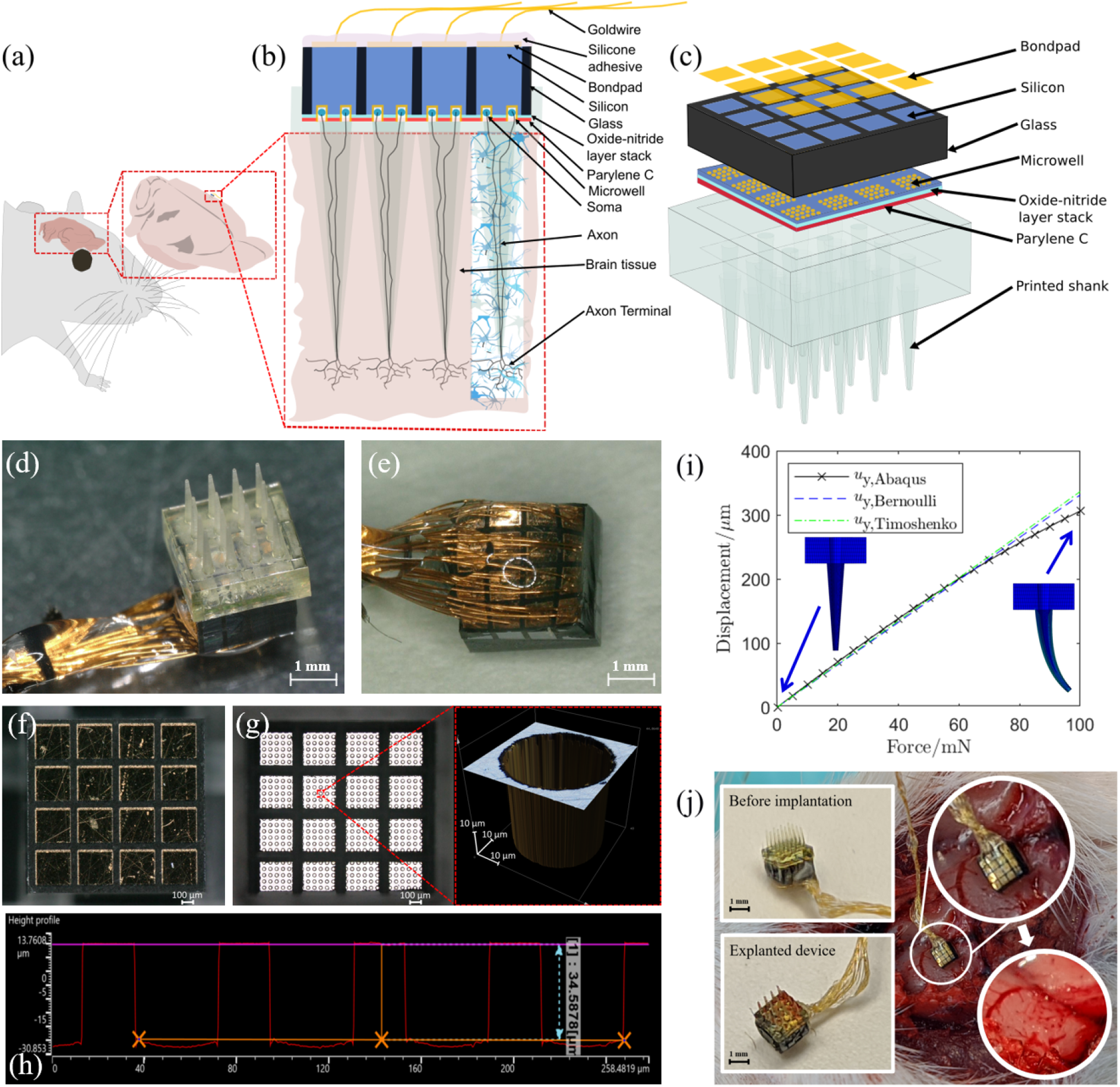
**a**, Schematic of the functional principle of the biohybrid TMEA, which is implanted into a rodent’s brain. Neurons that are trapped in a silicon-based chip’s micromachined golden wells project their neurites through hydrogel-filled microchannels of 4×4 polymer shanks towards the brain tissue. **b**, At the interface between the neuron culturing hydrogel and the brain tissue the tissue-engineered neural networks connect to the brain’s neural network through synapsis. This connection to the brain’s local neuron network is shown for one of the four shanks. **c**, Illustration of the biohybrid TMEA by means of an exploded view with a 4×4 array of electrode forming 3D-polymerized shanks. **d**, Actual realization of the biohybrid TMEA consisting of the silicon-based chip with µWells and the polymer shanks and **e**, backside view showing the bond wires for electrical connections of the 16 electrode sites as well as the silicone encapsulation. **f**, Image of the bondpads before bonding. **g**, Arrangement of the gold-coated µWells with a close-up of a single µWell measured with a laser microscope (Olympus Lext OLS5000).**h**, Depth profile and interspacing of the µWells. **i**, Investigation of the bending behavior of the 3D-polymerized shanks during implantation by means of a mechanical simulation. **j**, Implantation experiments using the pneumatic insertion technique into a dead rat’s brain to validate the polymer shanks’ mechanical stability.

Fig. 1 d and 1 e show the realization of the biohybrid TMEA neural implant (see also Supplementary Fig. S2). The device consists of two major parts: the hydrogel-filled polymer shanks and the silicon base chip. The conically shaped polymer shafts are created by advanced 3D lithography techniques (Two-photon polymerization, Nanoscribe Photonic Professional GT2) and measure 1 mm in length with a diameter of 150 µm at the base and 40 µm at the tip. The inner microchannel’s diameter is 20 µm, which can be modified as needed for different applications. As a neuron-growth enabling hydrogel material gelatin methacrylate (GelMA) was used. The channel voids were filled with the GelMA pregel solution by a vacuum-assisted suction technique. The pregel was subsequently UV-crosslinked to form the hydrogel (see Methods: *Hydrogel synthesis*). The silicon base shown in Fig. 1 d was fabricated using silicon micromachining techniques, which were derived from the fabrication process of the Utah Electrode Array [15] (see Methods: *Chip fabrication* and Supplementary Fig. S1 a). The base chip was designed to have 4×4 glass-isolated recording and stimulating sites on a 0.8 mm thick, 1.6 cm^2^ silicon substrate. Each site was equipped with a gold bondpad on the backside for electrical connection. The front side of the silicon base featured a configuration of 5×5 µWells at each recording site, yielding a total of 400 µWells on the device. Each of these µWells was gold-coated and had a lateral size and depth of 30 µm, in order to match the size of neuronal cells (see Fig. 1 f to 1 h). After the fabrication, murine cortical neurons prepared from pre-natal (E18) C57BL/6 mice (Jackson Labs, ME, USA) were extracted and pipette-loaded into the base µWells after fabrication of the silicon base (see Methods: *Neuron integration*). As a last step, the hydrogel-filled polymer shaft array and the neuron-loaded base chip were assembled then fixed to each other with a silicone adhesive (MED-4211, NuSil, Carpenteria, CA), thus completing the implant (see Supplementary Fig. S1 b). Two gold wires were wire-bonded to each of the 16 bondpads and sealed with silicone adhesive to encapsulate and mechanically reinforce the connection.

The polymer shafts are designed to be strong enough to penetrate the meninges surrounding the brain, survive the pneumatic insertion into the brain tissue and serve as a guidance channel to direct outgrowth into brain tissue. As the polymer shafts may be subject to mechanical failure by bending due to buckling during implantation, the shaft’s insertion behavior was investigated analytically and experimentally. A simulation of the mechanical behavior by means of a 3D Finite-Element model showed a bending stiffness of 280 to 325 N m^*−*1^, which is comparable to the stiffness of implantable optical fiber microelectrodes [16] (Fig. 1 i and Supplementary Fig. S3). A rat cadaver implantation experiment, where all shafts remained intact, confirmed these theoretical findings and demonstrated the device could be implanted into rat brain tissue without breaking (Fig. 1 j).

*In-vitro* accelerated aging tests with three biohybrid implants were performed to estimate the silicon and metal-based recordings sites’ lifetime. The accelerated aging experiments were carried out without hydrogel fillings and without neurons. The devices were kept at 67 ^*°*^C in phosphate buffered solution (1xPBS) for 46 days, which according to the Arrhenius equation corresponds to 1 year in tissue at a temperature of 37 ^*°*^C. Electrochemical impedance spectroscopy and cyclic voltammetry measurements were performed at various time intervals (see Methods: *Electrochemical stability*). With an average impedance amplitude reduction from 16.5 kΩ (±2.5 kΩ standard deviation) to 15.5 kΩ (±2.1 kΩ standard deviation) for 1 kHz the impedance and phase spectra did not show significant changes during the equivalent year of the study (see Fig. 2 a to 2 d). In addition, no mechanical-wire bonding disconnections or gradual material damage were observed. Furthermore, cyclic voltammetry tests confirmed the recording sites’ repeated charge storage stability, where median charge storage capacity for the illustrated device remained between 10^*−*5^ and 10^*−*6^ C cm^*−*2^ during the study period. These results imply that the implant body with its recording sites is viable for a one-year *in-vitro* or *in-vivo* recording study employing neurons.

**Fig. 2.**
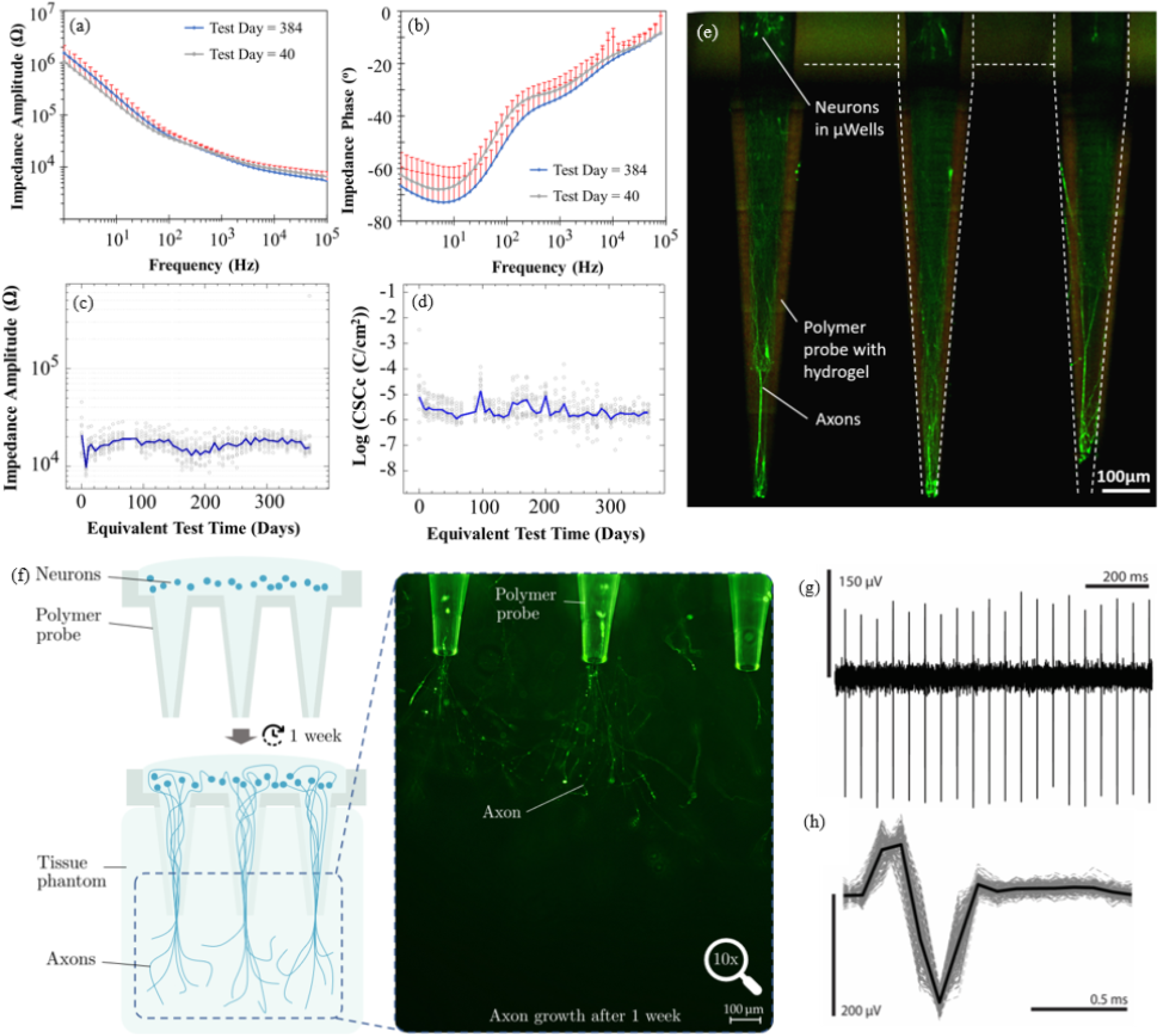
Accelerated aging measurements for our 16-channel biohybrid TMEA by means of a, impedance spectra and **b**, phase spectra over the course of 384 equivalent days (red bars represent the standard deviation, three devices were tested, one example device is shown in the graph). Time resolved **c**, median impedance and **d**, median charge-storage-capacity values at 1 kHz, respectively, for the same device (grey data points present individual electrode impedance values). **e**, A fluorescent confocal image of the Calcein-AM stained neurons reveals their neurites’ projections through the implant’s polymer shafts. **f** Axon growth through the shanks into a hydrogel-based tissue model. **g**,**h** show representative continuous time recording data from the generated 2 kHz sine wave spike trains and *in-vitro* recordings, respectively.

The formation of microtissue engineered neural networks within the polymer shanks was investigated by a fluorescent confocal image of calcein-AM stained neurons (see Fig. 2 e). For this experiment the extracted neurons were directly mixed into the GelMA pregel solution and immobilized within the hydrogel itself during the UV-initiated crosslinking step. The silicon base chip was not used for this experiment and the cells were filled in the polymer part directly. Projections occured successfully throughout the entire hydrogel bulk within the shafts’ guidance channels. Fig. 2 f demonstrates axon growth from neuron stem cells located at the base of the polymer probes through the shaft into a GelMA tissue phantom (see Methods: *Hydrogel synthesis*). Within a week after cell application fluorescence microscopic imaging of the stained axons revealed their successful projection. As a preliminary assessment of the biohybrid TMEA’s neuron recording capabilities, we evaluated its ability to record 2 kHz sinusoids, spaced 40 ms apart, as artificial neuronal signals in PBS solution (see Fig. 2 g and h). This experiment was performed with an implant that didn’t contain neurons and hydrogel, i.e. the shanks were left empty. The signals were introduced into the solution by playing them as audio files through a computer with its audio output connected to a Pt wire in the solution. The recordings were performed using the CerePlex Direct recording system and the CerePlex headstage (Blackrock Neurotech Inc.) with an analog bandpass filter from 250 Hz to 5 kHz and a 10 kHz sample rate (further details see Methods: *Neural signal recording*).

In this work, we presented a novel type of intracortical neural implant that uses a bioartificial construct with primary neurons instead of anorganic electrodes. Feasability aspects of the basic concept were demonstrated and included the fabrication of individual components which were subsequently assembled into a prospective brain implant. Insertion experiments into a rat’s brain and simulations showed that these microelectrodes of biohybrid TMEA withstand implantation without mechanical damage. Accelerated aging experiments and recording of artificial neural signals with unloaded devices indicated the recording circuitry’s functionality. Neuron-loaded devices proved a directed neuron outgrowth through the polymeric channels.

## Conclusion

Our current research efforts are on *in-vitro* signal recording with neuron-loaded devices. The study will be complemented by subsequent *in-vivo* trials conducted using rodent models. These experiments will involve readout over several weeks and subsequent immunohistochemical evaluation of brain slices to gain a better understanding of biocompatibility and connectivity with the host brain’s neural network. We anticipate that this biohybrid approach, employing endogenous and body-like materials, will result in a better human body implant acceptance and a comparatively long operating lifetime. Although the device’s polymer shaft material is still stiff, this can be replaced in the future by a soft or even biodegradable material, so that in the long term only the neuronal connection remains in the brain. Furthermore, we expect that with the use of biohybrid electrodes based on actual neuronal connections, an unprecedented spatiotemporal resolution may be provided. A single neuron is much smaller in dimensions than electrodes used to date, thus dramatically increasing the potential number of readout points per area. Our biohybrid TMEA approach would thus have great potential to significantly expand our current knowledge of brain processes.

## Supplementary Materials

Any methods, additional references, supplementary information, acknowledgements, details of author contributions and competing interests; and statements of data and code availability are provided in supplementary materials.

## References

[1] P. K. Campbell, K. E. Jones, R. J. Huber, K. W. Horch, and R. A. Normann. A silicon-based, three-dimensional neural interface: manufacturing processes for an intracortical electrode array. IEEE Transactions on Biomedical Engineering, 38(8):758–768, 1991.

[2] K. D. Wise, J. B. Angell, and A. Starr. An integrated-circuit approach to extracellular microelectrodes. IEEE Transactions on Biomedical Engineering, (3):238–247, 1970.

[3] P. R. Troyk and S. F. Cogan. Sensory neural prostheses. Neural Engineering, pages 1–48, 2005.

[4] M. Leber, J. Körner, C. F. Reiche, M. Yin, R. Bhandari, R. Franklin, S. Negi, and F. Solzbacher. Advances in penetrating multichannel microelectrodes based on the utah array platform. Neural Interface: Frontiers and Applications, pages 1–40, 2019.

[5] J. Leach, A. K. H. Achyuta, and S. K. Murthy. Bridging the divide between neuroprosthetic design, tissue engineering and neurobiology. Frontiers in Neuroengineering, page 18, 2010.

[6] T. Lim, M. Kim, A. Akbarian, J. Kim, P. A. Tresco, and H. Zhang. Conductive polymer enabled biostable liquid metal electrodes for bioelectronic applications. Advanced Healthcare Materials, 11(11):2102382, 2022.

[7] A. Ersen, S. Elkabes, D. S. Freedman, and M. Sahin. Chronic tissue response to untethered microelectrode implants in the rat brain and spinal cord. Journal of Neural Engineering, 12(1):016019, 2015.

[8] S. Musallam, M. J. Bak, P. R. Troyk, and R. A. Andersen. A floating metal microelectrode array for chronic implantation. Journal of Neuroscience Methods, 160(1):122–127, 2007.

[9] C. Malherbe, M. De Gasparo, R. De Hertogh, and J. J. Hoet. Circadian variations of blood sugar and plasma insulin levels in man. Diabetologia, 5(6):397–404, 1969.

[10] J.M. Hsu, L. Rieth, R. A. Normann, P. Tathireddy, and F. Solzbacher. Encapsulation of an integrated neural interface device with parylene c. IEEE Transactions on Biomedical Engineering, 56(1):23–29, 2008.

[11] P. J. Schubert and J. H. Nevin. A polyimide-based capacitive humidity sensor. IEEE Transactions on Electron Devices, 32(7):1220–1223, 1985.

[12] D. Vega, J. Reina, F. Martí, R. Pavón, and Á. Rodríguez. Macroporous silicon for highcapacitance devices using metal electrodes. Nanoscale Research Letters, 9(1):1–8, 2014.

[13] C. H. Chiang, C. Wang, K. Barth, S. Rahimpour, M. Trumpis, S. Duraivel, I. Rachinskiy, A. Dubey, K. E. Wingel, M. Wong, N. S. Witham, T. Odell, V. Woods, B. Bent, W. Doyle, D. Friedman, E. Bihler, C. F. Reiche, D. G. Southwell, M. M. Haglund, A. H. Friedman, S. P. Lad, S. Devore, O. Devinsky, F. Solzbacher, B. Pesaran, G. Cogan, and J. Viventi. Flexible, highresolution thin-film electrodes for human and animal neural research. Journal of Neural Engineering, 18(4):045009, 2021.

[14] D. O. Adewole, L. A. Struzyna, J. C. Burrell, J. P. Harris, A. D. Nemes, D. Petrov, R. H. Kraft, H. I. Chen, M. D. Serruya, J. A. Wolf, and D. K. Cullen. Development of optically controlled living electrodes with long-projecting axon tracts for a synaptic brain-machine interface. Science Advances, 7(4):eaay5347, 2021.

[15] R. Bhandari, S. Negi, and F. Solzbacher. Wafer-scale fabrication of penetrating neural microelectrode arrays. Biomedical Microdevices, 12:797–807, 2010.

[16] S. Park, H. Yuk, R. Zhao, Yeong S. Yim, E. W. Woldeghebriel, J. Kang, A. Canales, Y. Fink, G. B. Choi, X. Zhao, and P. Anikeeva. Adaptive and multifunctional hydrogel hybrid probes for long-term sensing and modulation of neural activity. Nature Communications, 12(1):1–12, 2021.

