## Supplementary Materials for "Implementation and testing of a biohybrid transition microelectrode array for neural recording and modulation"

#### This PDF file includes:

Methods
Figs. S1 to S3
Tables S1 to S6

### Methods

**Chip fabrication.** The fabrication process for the biohybrid microelectrode array is illustrated in supplementary materials. Fig. S1. Specifications of the used Silicon wafer are given in Table S1. The fabrication process is in parts based on the Utah Electrode Array fabrication.

*Recording site creation* - First, to create the trenches for the glass insulation of the recording sites, a dicing saw (DAD 3220, Disco, Japan) was used on the polished side of the the wafer. In a batch process, for each device five cuts were made in each horizontal and vertical direction, respectively (depth 800 µm, pitch 400 µm). This results in arrays of 4x4 silicon islands, which will later be the recording sites. In total, cuts for 200 devices were carried out on a single wafer. After dicing, the kerfs were cleaned in ultrasonic baths of isopropyl alcohol (IPA) and subsequently deionized (DI) water to remove silicon debris followed by drying on a hotplate at 150 °C for 5 min. Fig. S1 a (i) and (ii) show schematics of the backside dicing process and the wafer after backside dicing, respectively.

*Recording site isolation* - Glass frit (EG 2790 VWG, Ferro Corporation, USA) was mixed into methanol (1:1 by wt.) and applied to the grooved surface and then dried in an oven at 60 °C for 5 min. The silicon wafer was then placed into a ceramic boat and loaded into a dental vacuum furnace (Ney Centurion Qex, Denstply Ceramco, USA). The furnace temperature was gradually raised to 1150 °C at a rate of 25 °C/min, facilitating the melting of the glass, which effectively filled the kerfs. By turning off the furnace, the temperature was decreased back down to room temperature slowly solidifying the liquid glass in the kerfs. The glassing procedure left an uneven layer of excess glass on the wafer (see Fig. S1 a (iii)), which was removed by a grinding-polishing process. This grinding-polishing process was done by removing the bulk glass from the surface with dicing saw (DAD 3220, Disco, Japan), which stripped off excess glass, revealing an uneven silicon surface with residual glass fragments (see Fig. S1 a (iv)). Subsequently, the wafer was manually polished with Silicon Carbide (SiC) abrasive papers (1200 grit/P2500, Buehler, USA) for approx. 60 min. The polishing procedure involved a 3 µm polycrystalline diamond suspension (40-6631, Metadi Supreme™, Buehler, USA) for the first 30 min of the process, then a 1 µm polycrystalline diamond suspensions (40-6630, Metadi Supreme™, Buehler, USA) for the remaining 30 min, which resulted in a crosshatch pattern of insulating glass as shown in Fig. S1 a (v). At this process stage, the silicon islands (recording sites) were electrically still connected by the silicon on the wafer's opposite side. To electrically isolate these recording sites, the wafer was flipped over and subsequently grinded and polished as described in the previous de-glassing process. This resulted in 4x4 arrays of silicon islands, i.e., the recording sites, which were physically connected by glass lines, while at the same time, electrically isolated from each other.

*Bondpad creation* - Metal bondpads were created on the back side of each recording site (see Fig. S1 a (vi)) by performing a lift-off process. This was achieved through photolithographic structuring of a photoresist (S1813, Shipley Company Inc., USA) with a UV exposure dose of 60 mJ/cm$^2$), followed by DC sputter-depositing a metal stack consisting of 300 nm Gold, 100 nm Platinum and 50 nm Titanium (Au/Pt/Ti) using a sputtering machine (SS-40C-IV, TM Vacuum Products, USA). The lift-off was subsequently done by submerging the wafer in an ultrasonic acetone bath resulting in 320×320 µm$^2$ Au/Pt/Ti contact pad on each of the glass-isolated recording sites. The bonds were able to pass a scotch tape test and, hence, exhibited good adhesion with the silicon substrate.

*µWell fabrication* - Four 250 nm thick layers of alternating silicon oxide and silicon nitride (total thickness: 1 µm) were deposited on the front side of the wafer via plasma enhanced chemical vapor deposition (Plasmalab 80 Plus, Oxford Instruments Plasma Technology, UK) to improve the electrical isolation between the adjacent neural recording sites (see Fig. S1 a (vii). The process parameters are provided in Table S2 and S3. To improve the biocompatibility of the recording sites, a 6 µm thick parylene C film was deposited using a low temperature (25 °C) chemical vapor deposition process with a parylene C caoter (PDS 2010, Specialty Coating Systems, USA) while the backside of the wafer containing the bondpads was protected with tape to prevent parylene C from settling on the bondpads. As an adhesion promoter (Silquest A-174 silane, GE Silicones Inc., USA) was used. To create the µWells, i.e., the neuron culture sites, the parylene C, oxide-nitride layer stack and the silicon underneath three separate etching steps were required to separately etch parylene C, oxide-nitride layer stack, and silicon (see Fig. S1 a (viii)-(xii)). The etching mask was carefully fabricated through a photolithographic process utilizing a photoresist (AZ 9260, AZ Electronic Materials, Luxembourg) with a UV exposure dose of 400 $mJ/cm^2$ , along with a developer (AZ 400K, AZ Electronic Materials, Luxembourg). Subsequently, parylene C was plasma-etched (Oxford Plasmalab 100, Oxford Instruments Plasma Technology, UK). Please see Table S4) and Table S5) for parylene C and oxide-nitride layer stack etching parameters respectively. Finally, the silicon was etched using the Bosch process, i.e. deep reactive ion etching, by alternating between etching using a $SF_6$ plasma pulse and creating a protective layer using a $C_4F_8$ plasma pulse. The $SF_6$-$C_4F_8$ cycle was repeated 20 times to reach an overall depth of 35 µm (see Table S6 for process parameters). To facilitate neuronal cell growth and the recording of neural signals, the walls of the µWells were gold-coated. This was done by microstructuring an Au/Pt/Ti metal stack (total thickness: 450 µm) by means of a lift-off process as described previously (see Fig. S1 a (xiii-xiv)). Prior to the metal sputtering into the µWells, the $C_4F_8$ passivation layer from the Bosch process was removed with oxygen plasma cleaning (PlanarEtch IIA plasma system,Technics Inc., USA) using DC power 100 W for 3 minutes to allow deposition of metal layers directly on silicon walls of the µWells. Finally, the photoresist layer was lifted off using acetone and the wafer was then put through an ultrasonic cleaning process using IPA, followed by a rinse with deionized water same as before. The metallized µWells on the silicon base chip are shown in Fig. S2 a.

*Device singulation and wirebonding -* Finally, each of the silicon base chips containing 4x4 neural recording sites was singulated from the wafer using dicing machine(Disco DAD 3220, Disco, Japan). After the individual base chips had been separated, they were all subjected to ultrasonic cleaning with IPA and DI water. A semi-automatic wire bonder (545657E, West Bond Inc., USA) was used to attach insulated Au wires of 56 µm diameter (PF4418, 1% Pd, Treseter966, Sandvik Palm Coast, USA) to the metallic bondpads (refer to Fig. S1 a (xv)). The other end of these bond wires were wire-bonded to a connector for the neural recording instrument, thus enabling electrical contacts between the neural recording sites and the neural recording instrument. The wirebundle was reinforced by using a biocompatible silicone adhesive (MED-4211, NuSil, USA) as shown in Fig. S2 b for mechanical and electrical reinforcement.

**Polymer shank fabrication.** The polymer shanks were designed to measure 1000 µm in length and have a 700 µm base with a rectangular cavity (1600 µm x 1600 µm in length and width, and 400 µm in depth), allowing for integration with the silicon base chip. The pitch between the shanks was designed to be 400 µm to match the silicon base chip's recording site pitch, making it possible to align each shank with an individual neural recording site. The diameters of the shank's tip and base were designed to be 40 µm and 100 µm, respectively, while the wall thickness of the hollow shank was 20 µm. These polymer shanks were 3D-polymerized with a two-photon polymerization process (Nanoscribe Photonic Professional GT2, Nanoscribe GmbH, Germany) using IP-S resin, thus forming shanks with integrated microchannels to guide projected neurites from neuronal cells to a particular direction. Fig. S2 c shows an image of the 3D-polymerized shanks.

**Hydrogel synthesis.** Gelatin methacrylate (GelMA) hydrogel is a photo-polymerizable methacrylated gelatin with excellent biocompatibility [1]. Gelatin can be engineered to become photo-crosslinkable via the controlled incorporation of methacrylate groups to the amine-containing side groups, which enables the fabrication of complex geometries. Additionally, GelMA is an attractive candidate for rapid in-situ photopolymerization. In this study, we synthesized GelMA based on previously reported protocols [2, 3]. Gelatin (type A, 300 bloom from porcine skin), methacrylic anhydride (MA), 3-(trimethoxysilyl)propyl methacrylate (TMSPMA) and the photoinitiator Lithium phenyl-2,4,6-trimethylbenzoylphosphinate (LAP) were purchased from Sigma-Aldrich, USA. The synthesized GelMA was suspended in Dulbecco's phosphate buffered saline (DPBS, 5%, w/v) and mixed with 0.5% (w/v) LAP. The pregel was exposed to

UV light for 10 to 12 s at a power density of 3.46 mW cm$^{-2}$ to initiate the crosslinking reaction resulting in a hydrogel.

**Neuron integration.** The isolation of the murine cortical neural progenitor cells was conducted in compliance with the guidance and policies of the Institutional Animal Care and Use Committee (IACUC). Primary neuron cultures were prepared from male and female E18 murine cortex as previously described [4]. Tissue was dissociated in 20U/mL papain (Worthington Biochemical Corp) and DNase (Sigma), and then triturated with fire-polished glass pipettes to obtain a single-cell suspension. Cells were pelleted at 500×g for 5 min, the supernatant removed, and cells resuspended in Neurobasal media containing 5% fetal bovine serum, 1% glutamAX, 1% penicillin/streptomycin (Thermo Fisher Scientific), and 2% SM1 (Stem Cell Technologies).
Dissociated neuronal cells were pipette-loaded to the µWells of the biohybrid microelectrode arrays. The loaded neurons were trapped in the µWells. Since the dimensions of µWells are designed to hold only one single cell, each µWell can be filled by one single neuron after several cell loading-washing cycles. Extra cells were washed away. This process is schematically shown in Fig. S1 c.

**Device assembly.** The 3D-polymerized and hydrogel-filled polymer shaft array was assembled with the neuron-loaded silicon base chip to get the complete brain implant device (see Fig. S1 d). A silicone adhesive (MED-4211, NuSil, USA) was used to fix those two components to each other. Fig. S2 d shows an image of the complete biohybrid brain implant device after assembly. The entire device was put into a cell incubator (Forma 320, Thermo Fisher Scientific, USA) for *in-vitro* cell culture.

**Mechanical simulation.** Multi-field modelling and simulation can be applied as a tool to design the shanks such that the piercing process leads to minimal damaging of the brain tissue and the grown neurites inside the shanks are mechanically protected, while the diffusion of growth factors to the neurons is guaranteed. In the current work, the simulation focuses on describing mechanical reactions that occur during the implantation process, i.e., (A) bending and (B) buckling.

(A) The polymer shafts (1 mm long, 150 µm at the bottom and 40um on the top) need to be strong enough to penetrate the cortical surface and survive the pneumatic insertion into the brain tissue and may be subject to mechanical failure and bending during this process. The simulations were performed in Abaqus as a half-model due to symmetry and loading. The material is IP-S (Neo-Hookian material model with C10 = 850 MPa and D1 = 1.1765e-05) and the geometry was extracted from the CAD-model for 3D-polymerization. A total of 7596 quadratic hexahedral elements of type C3D20H were used for the simulation.

(B) Static buckling instability is a mechanical reaction that occurs for slender beam-like structures under pressure loading. At a critical load $F_{\mathrm{critical}}$, multiple force-displacement paths can occur (bifurcation). For simple beam structures and boundary conditions, the Euler-cases can be derived from the differential equations for the deformed state of the beam [5]. Buckling can happen in the slim shanks of the device structure. When a shank is inserted into brain tissue, the maximum force which is relevant for the buckling can be found either (i) directly at the initial piercing of the brain tissue by the shaft or (ii) when the shaft is already inserted, and a higher force is needed with increasing piercing depth, see Fig. S3 a. For the relevant Euler cases, the critical length is defined (i) for the initial piercing as $l_{\mathrm{critical,(i)}} = 0.7 \cdot 1000\,\mu\mathrm{m}$ and (ii) for the further insertion as $l_{\mathrm{critical,(ii)}} = 0.5 \cdot (1000\,\mu\mathrm{m} - d_{\mathrm{Piercing}})$, whereas the insertion depth is $d_{\mathrm{Piercing}}$. The radius of gyration is $\sqrt{I(s)/A(s)}$ with the moment of inertia $I(s)$ and the cross-section area $A(s)$ which are both dependent on the position $s$, due to the conical shape of the shanks. To provide a conservative assessment of the slenderness, the minimum radius for $s$=0 is considered. The according slenderness factors are $\lambda_{((i))} \approx 39$ and $\lambda_{((ii))} \approx 28$ (for $d_{\mathrm{Piercing}} = 0\,\mu\mathrm{m}$) which are both much higher than 17, which means that the structures are prone to buckling [5]. The critical buckling forces can be calculated by

$$F_k = \pi^2 \frac{EI}{l^2_{critical}} \tag{1}$$

which leads to $F_{(\mathrm{k,(i)})} = 52.4\,\mathrm{mN}$ and $F_{(\mathrm{k,(ii)})} = 102.8\,\mathrm{mN}$. These simplified calculations provide an insight into the order of magnitude in which the critical forces can be found. Due to the complexity of both geometry and constraints in the tMEA structure, Finite-Element simulations of the structure were performed to gain the critical loads. To obtain a Finite-Element solution of the critical load $F_{\text{critical}}$, the system behavior is linearized at the critical point. A load increment is defined, which leads to an eigenvalue problem, the solutions are critical loads (eigenvalues) and buckling mode shapes (eigenvectors). In the current work, we apply the subspace method in Abaqus to solve the eigenvalue problem. The Finite-Element results of Abaqus leads to buckling shapes and critical loads as shown in Fig. S3 a. In comparison to the simplified analytical derivations with the Euler cases, the critical load is higher by the factor 5 for case (i) and the factor 4.5 for case (ii). This is due to the simplification of the constant wall thickness. In Fig. S3 b, the increasing critical load with increasing piercing depth is depicted. It is found that after the initial insertion, the structure gets less prone to buckling due to the much higher increase of critical load compared to the expected increasing of the reaction force due to wall friction.

**Electrochemical stability.** Each of the three devices was kept in 1xPBS baths of 67 °C to reach an aging acceleration factor of eight according to an Arrhenius aging assumption for polymeric materials [6]. The arrays were aged for 46 days equivalent to approx. one year of aging in a human body environment. Except for short testing periods, samples were kept inside the hot 1xPBS bath. Electrochemical impedance spectroscopy (EIS) and cyclic voltammetry (CV) measurements were conducted with a potentiostat (Reference 600, Gamry Instruments, USA). A frequency range of 1 Hz to 100 kHz with an AC signal of 50 mV versus open circuit potential was used for EIS. An additional pair of platinum wire were used as reference/counter electrodes. For CV, voltammograms were recorded at 50 mV ms$^{-1}$ slope in the water electrochemical window from -0.6 V to 0.8 V. Three cycles of 50 50 mV s$^{-1}$ CVs were recorded at each test section for each electrode to reach stability. The electrochemical measurements were carried out using a custom-built 96-channel automatic switch board. All EIS and CV data were recorded daily for every individual electrode. The CV results were used to calculate the charge storage capacity (CSC) of each electrode and the electrode's potentiodynamics.

The data analysis was carried out in MATLAB, Excel and Origin software packages.

**Neural signal recording.** As a preliminary assessment of the biohybrid brain implant's recording site, we evaluated an empty (i.e. without hydrogel and neurons) device's ability to record generated signals meant to approximate the shape and frequency composition of neuronal action potentials. These artificial signals consisted of individual 2 kHz sinusoids spaced approximately 40 ms apart. To facilitate the signal transmission, the generated signal was converted to a sound file and played through a severed earbud cable, connected to the audio jack of the computer, with a single signal and ground wire exposed. The signal wire was connected to a platinum wire in solution with the biohybrid brain implant and the ground wire was connected to the ground and reference of the recording system. All recordings were made using the CerePlex Direct recording system and Cereplex headstage (Blackrock Neurotech, USA) with an analog bandpass filter from 250 Hz and 5 kHz and a 10 kHz sampling frequency. The signal-to-noise-ratio calculations for the mean waveform of the threshold crossing events were made using the method described in [7, 8]. Briefly, the peak-to-peak amplitude of the mean waveform $\bar{W}$ is divided by the standard deviation of the noise in the waveform

$$SNR = \frac{\max(\bar{W}) - \min(\bar{W})}{SD_\epsilon}, \qquad (2)$$

where $\epsilon$ is a matrix of the difference of each point of each individual waveform from the mean.

### Data availability

All raw data generated during this project are available from the authors upon request.

### References

[1] A. I. Van Den Bulcke, B. Bogdanov, N. De Rooze, E. H. Schacht, M. Cornelissen, and H. Berghmans. Structural and rheological properties of methacrylamide modified gelatin

hydrogels. *Biomacromolecules*, 1(1):31–38, 2000.

[2] J. W. Nichol, S. T. Koshy, H. Bae, C. M. Hwang, S. Yamanlar, and A. Khademhosseini. Cell-laden microengineered gelatin methacrylate hydrogels. *Biomaterials*, 31(21):5536–5544, 2010.

[3] Y. Fan, F. Xu, G. Huang, T. J. Lu, and W. Xing. Single neuron capture and axonal development in three-dimensional microscale hydrogels. 12(22):4724.

[4] E. D. Pastuzyn, C. E. Day, R. B. Kearns, M. Kyrke-Smith, A. V. Taibi, J. McCormick, N. Yoder, D. M. Belnap, S. Erlendsson, D. R. Morado, and J.A. Briggs. The neuronal gene arc encodes a repurposed retrotransposon gag protein that mediates intercellular rna transfer. *Cell*, 172(1):275–288, 2018.

[5] C. Spura. *Einführung in die Balkentheorie nach Timoshenko und Euler-Bernoulli*. Springer, 2019.

[6] D. W. L. Hukins, A. Mahomed, and S. N. Kukureka. Accelerated aging for testing polymeric biomaterials and medical devices. *Medical Engineering & Physics*, 30(10):1270–1274, 2008.

[7] R. C. Kelly, M. A. Smith, J. M. Samonds, A. Kohn, A. B. Bonds, J. A. Movshon, and T. S. Lee. Comparison of recordings from microelectrode arrays and single electrodes in the visual cortex. *Journal of Neuroscience*, 27(2):261–264, 2007.

[8] M. Sharma, A. T. Gardner, H. J. Strathman, D. J. Warren, J. Silver, and R. M. Walker. Acquisition of neural action potentials using rapid multiplexing directly at the electrodes. *Micromachines*, 9(10):477, 2018.

#### Acknowledgements

This research is based upon work supported by the National Institutes of Health (NIH) under Brain Initiative Award No 1R21EY033082-01. The content is solely the responsibility of the authors and does not necessarily represent the official views of the NIH.
Simon Binder acknowledges funding by the Deutsche Forschungsgemeinschaft (DFG, German Research Foundation) – 459675326.
This work was performed in part at the Utah Nanofab sponsored by the University of Utah's College of Engineering, Office of the Vice President for Research, and the Utah Science Technology and Research (USTAR) initiative of the State of Utah. The authors appreciate the support of the staff and facilities that made this work possible. Patrick Tresco's and Austin Koch's consultation and assistance with experiments gratefully acknowledged.

#### Author contributions

Contributions according to CRediT:

- Conceptualization (Y.F., F.S., St.B., C.F.R.)
- Methodology (Si.B., M.H., A.E.)
- Investigation, validation, formal analysis (B.A., Si.B., Sa.B., H.J.S., M.H., T.S., A.K., A.E.)
- Writing original draft (B.A., Si.B., Sa.B., H.J.S., M.H., T.S., A.K., A.E.)
- Review & editing (Y.F., St.B., F.S., P.A.T., H.Z., C.F.R.)
- Supervision (Y.F., St.B., F.S., P.A.T., C.F.R., J.S., H.Z.)
- Funding acquisition (Y.F., St.B., C.F.R., F.S.)
- Project administration (Y.F., St.B., C.F.R., F.S.)

#### Competing interests

The authors declare the following competing financial interests that are overseen through the University of Utah conflict of interest management:
Fan, Y., Reiche, C. F. and Solzbacher, F. *Implantable Transition Microylectrode Array*, 00846-U6846.PCT, 2022.
F. Solzbacher declares a financial interest in Sentiomed, Inc. and Blackrock Neurotech.

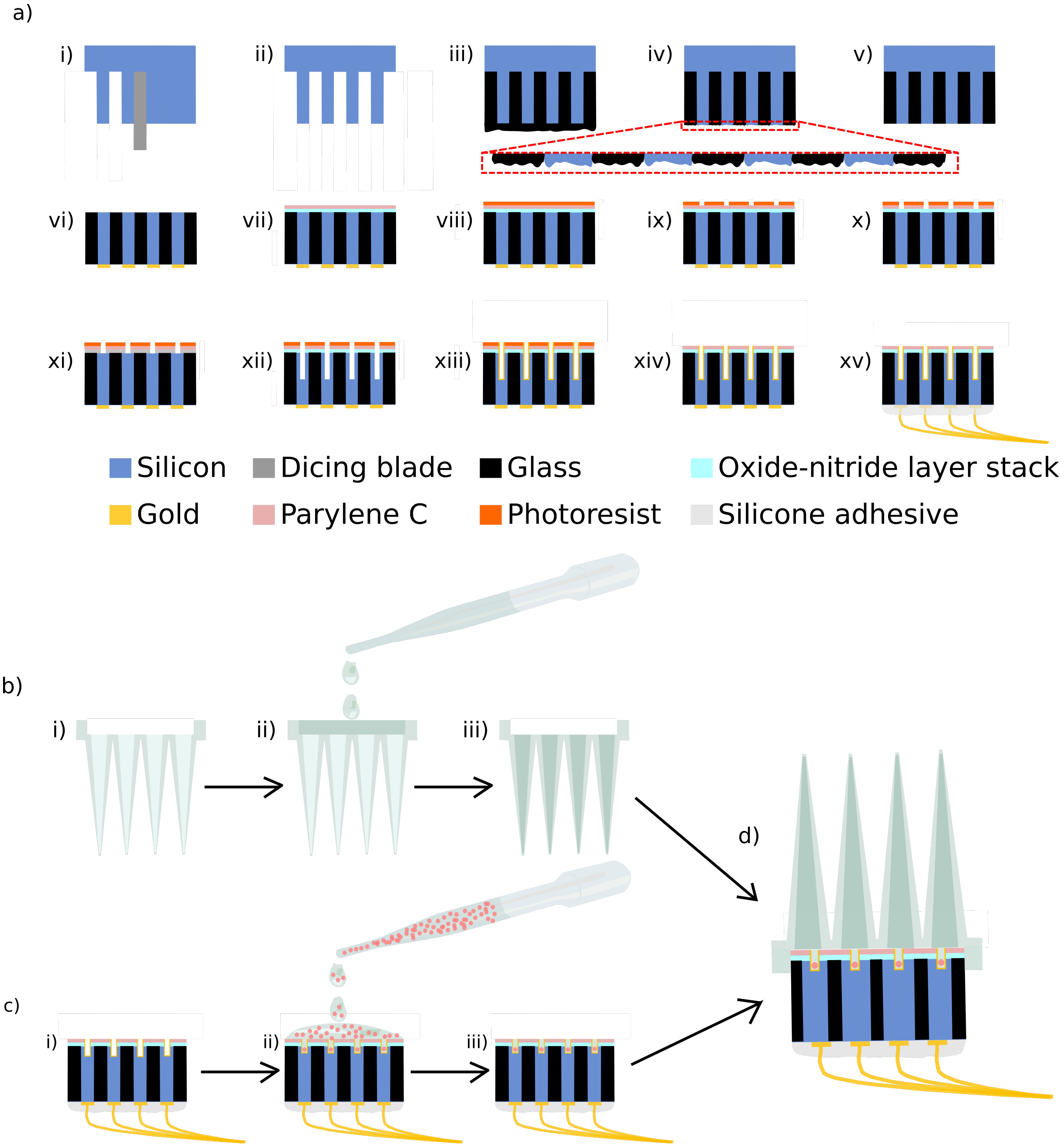


**Fig. S1** | **a,** Schematic of the fabrication steps of the silicon base chip: **(i)**backside dicing, **(ii)** silicon islands standing on silicon backplane after backside dicing, **(iii)** silicon kerfs filled with glass, **(iv)** silicon base chip after deglassing (inset indicated the rough surface), **(v)** silicon base chip with smooth surface after polishing, **(vi)** removal of silicon from the front side and bondpad metallization, **(vii)** oxide-nitride layer stack and parylene C deposition, **(viii)** deposition of photoresist, **(ix)** photolithography to create etching mask, **(x)** etching of parylene C, **(xi)** etching of oxide-nitride layer stack, **(xii)** etching of silicon to create µWells, **(xiii)** metal deposition on the µWells, **(xiv)** lift-off and **(xv)** wire bonding with insulated gold wires. **b,** GelMA loading in hollow shanks: **(i)** 3D-polymerized shanks with integrated microchannels, **(ii)** dispensing GelMA on the rectangular cavity with a pipette and **(iii)** suction induced loading of GelMA into the microchannels. **c,** Loading of neural cells in silicon base chip: **(i)** Silicon base chip, **(ii)** pipette loading of neural cells with culture medium on the base chip, **(iii)** cells in µWells after rinsing excess cells **d,** Biohybrid microelectrode array after integration of neural cell-loaded silicon base chip and GelMA loaded 3D-polymerized polymer shanks.

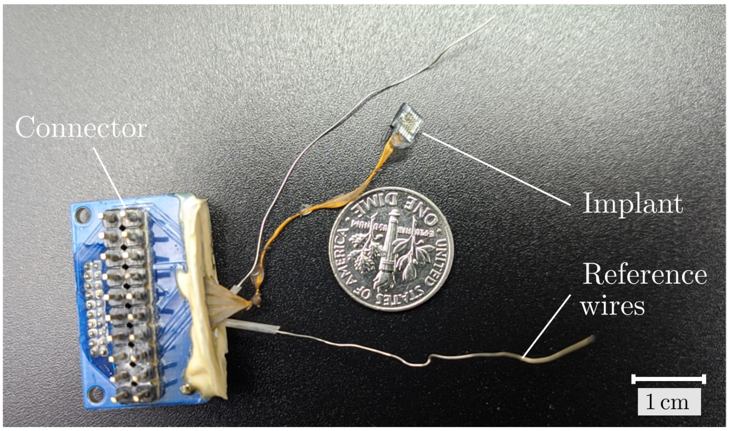


**Fig. S2** | Realization of the whole biohybrid neural implant consisting of the silicon base chip with the polymer shanks and the connector for the neural recording instrument including two reference electrodes.

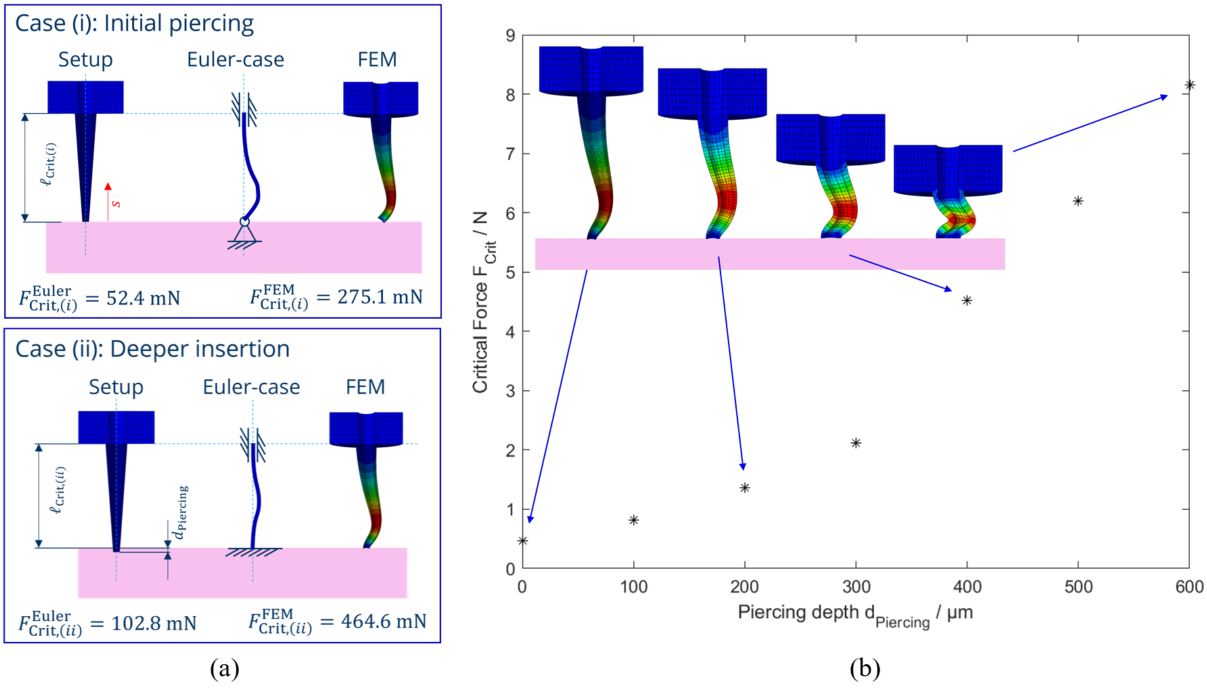


**Fig. S3** | **a,** Relevant buckling cases and equivalent beam model and buckling results for the first eigenmode of cases **(i)** and **(ii)** according to main text Fig. 1 (i) with a 100x deformation scale factor, **b,** piercing depth dependent critical force for Case (ii). The mesh is the same as for the bending simulation, a total of 7596 quadratic hexahedral elements of type C3D20H are used for the simulation. Boundary conditions are chosen according to the cases in (a).

**Table S1** Silicon wafer specifications

| | |
|---|---|
| Material | Silicon |
| Diameter | 76.2 mm ± 0.3 mm |
| Orientation | <100>±0.9° |
| Dopant | Boron |
| Resistivity | 0.01-0.04 Ω-cm |
| Center thickness | 2000 ±50 µm |
| Total thickness variation | <15 µm |

**Table S2** Oxide deposition parameters

| | |
|---|---|
| Chamber pressure | 1 Torr |
| $N_2$ flow | 161 sccm |
| $N_2O$ flow | 710 sccm |
| $SiH_4$ flow | 8.5 sccm |
| HF forward power | 21 W |
| Substrate temperature | 300 |

**Table S3** Nitride deposition parameters

| | |
|---|---|
| Chamber pressure | 1 Torr |
| $N_2$ flow | 380 sccm |
| $SiH_4$ flow | 20 sccm |
| $NH_3$ flow | 20 sccm |
| HF forward power | 21 W |
| LF power | 24 W |
| Substrate temperature | 300 |

**Table S4** parylene C etching parameters

| | |
|---|---|
| Chamber pressure | 25 mTorr |
| Helium back pressure | 10 mTorr |
| Oxygen flow | 50 sccm |
| RF forward power | 200 W |

**Table S5** Oxide-nitride layer stack etching parameters

| | |
|---|---|
| Chamber pressure | 10 mTorr |
| Helium back pressure | 10 mTorr |
| $CF_4$ flow | 50 sccm |
| RF forward power | 25 W |
| ICP forward power | 500 W |

**Table S6** Silicon etching parameters

| | |
|---|---|
| Helium back pressure | 10 mTorr |
| $SF_6$ flow | 80 sccm |
| $C_4F_8$ flow | 80 sccm |
| RF forward power | 25 W |
| ICP forward power | 500 W |
| $SF_6$ pulse time | 14 s |
| $C_4F_8$ pulse time | 5 s |
